# Biomass accumulation and physiological responses of tomato plants to magnetically–treated water in hydroponic conditions

**DOI:** 10.1101/2021.09.21.461287

**Authors:** Daniel I. Ospina-Salazar, Shimon Rachmilevitch, Santiago Cuervo-Jurado, Orlando Zúñiga-Escobar

## Abstract

Magnetically-treated water (MTW) has been reported to enhance biomass accumulation in plants. However, the underlying mechanisms are not fully understood, and the existing reports only deal with soil-grown plants. Thus, the purpose of this experiment was to assess whether or not MTW affects main physiological processes (gas exchange, biomass accumulation and water potential) in tomato plants whose water supply was only MTW. Two experiments were done in hydroponic semi-controlled conditions, consisting of a loop system with permanent recirculation of water through a non-uniform magnet. The plants grown under MTW showed a significant increase in chlorophyll content, photosynthesis and transpiration at high light irradiances, although the increase in stomatal conductance was less significant. MTW also increased fruit fresh biomass, number of fruits and root dry biomass in 61.7 %, 85.3 % and 30.3 % respectively, but this was only achieved at natural sunlight conditions. Moreover, treated plants showed higher root hydraulic conductance and leaf water potential, which is thought to be related with a lower surface tension of MTW, an effect that is consistent with previous studies. The higher biomass accumulation in tomato plants under MTW is likely explained because of a faster water transport from the roots to the leaves via xylem, which in turn increases H_2_O efflux and CO_2_ assimilation in the leaves, thanks to a higher stomatal conductance.

## Introduction

The physiological improvement of crops is the ultimate goal to achieve higher yields with lower usage of resources. Regarding this, implementation of magnetic treatments (either to the water or to the seeds) is an alternative that remains obscure in its basic physiological mechanisms. However, several breakthrough findings over the last years have unveiled important insights into specific processes, particularly in magnetoprimed seeds, such as enzymatic activity during germination (Radhakrishnan and Ranjitha-Kumari, 2012; Vashisth and Nagarajan, 2010a), seed water uptake (Bhardwaj et al., 2012; Vashisth and Nagarajan, 2010b) and photosynthetic activity in seedlings (Shine et al., 2011). As for the magnetic treatment of water, its physiological effects on plants are less known, but it has been reported that the activities of enzymes as catalase and peroxidase of tomato seedlings were increased when irrigated with magnetized water (Chen et al., 2008).

Although water, as well as living beings, is typically a diamagnetic substance (unlike ferromagnetic substances such as iron, nickel or cobalt), it can react to magnetic fields through modifications at a molecular level, not necessarily associated with an electron spin alignment. Such responses to magnetic fields involve different physicochemical properties, like surface tension, viscosity, evaporation rate and clustering of H_2_O molecules. At a molecular level, the work of Chang and Weng (2006) indicates that, under the influx of a magnetic field, there is a greater formation of hydrogen bonds between water molecules. The latter means that water clusters can be modified in its size depending on the applied magnetic induction. According to Toledo et al. (2008), magnetic treatment of water drives the formation of smaller water clusters but with stronger bonds. However, results from Cai et al. (2009) indicate that the MTW leads the formation of larger clusters, and this is related with a promotion in the activation energy. In a large review of hydrogen-bond relaxation dynamics, Huang et al. (2015) concluded that a magnetic field can mediate in the cooperative relaxation in the hydrogen bond. In any case, the experimental and theoretical data suggest that there is a competition between the intramolecular and intermolecular forces in water subjected to magnetic fields, resembling those that occur when there are changes in temperature.

The reduction in surface tension of magnetically treated water is an effect consistently observed under different experimental conditions (Lee et al., 2013; Huo et al., 2011; Cai et al., 2009; Amiri and Dadkhah, 2006; Otsuka and Ozeki, 2006). This effect is related with a decrease in hydrophobicity, evinced by the narrower contact angle of a sessile droplet on a smooth surface (Pang and Deng, 2008). Surface tension is a key factor that contributes to the ascent of sap along the xylem’s vessels and tracheid; several surfactant substances that have been detected in *Populus tremuloides* trees suggests that reduction of surface tension in the sap is important to reduce the risk of embolism (Christensen-Dalsgaard et al., 2011) and makes more hydrophilic the hydrophobic surfaces of the xylem (McCully et al., 2014).

Studies on magnetic treatment of water for crop irrigation have been conducted for decades (Volkonskii et al., 1978). Several authors have reported that MTW for crop irrigation leads to an increase in plant productivity and changes in its water and mineral metabolism (Hozayn et al., 2013; Al-Khazan et al., 2011; Maheshwari and Grewal, 2009). By using a specific magnetic device for water treatment, two authors (Grewal and Maheshwari, 2011; Maheshwari and Grewal, 2009) observed a general increase in the productivity of snow pea and chickpea based on fresh pod weight and shoot/root fresh and dry weight, respectively. The water productivity of snow peas also increased by 12 %, as well as the content of N, K, Ca, Mg, S, Zn, Fe and Mn in snow pea and chickpea seedlings. However, the authors make clear that while these findings are interesting, the potential of the MTW for crop production needs further testing to prove if whether it has any positive effect on the yield in relation to water use. Regarding the prior, irrigation with MTW increased the relative water content and water use efficiency in jojoba plants, at different moisture regimes, effect that was preserved even in the treatments at low moisture regimes, (Al-Khazan et al., 2011). This indicates that the ability of plants under low water regimes to manage water is optimized. Putti et al. (2015) and Hozayn et al. (2013) also reported an increase in water use efficiency and in green biomass in sugar beet and lettuce supplied with magnetically treated water, respectively. Despite the latter results, this technology is not widely used, in part due to the lack of a sound physiological background.

In order to contribute to the understanding of the mechanisms of MTW in plants, we tested the hypothesis that, in a controlled growing system where MTW is the only water source, this treatment affects physiological processes such as water transport, gas exchange and biomass accumulation, and that these physiological effects are related with the physicochemical changes of MTW. The experiment was conducted with a representative and short-cycle species, tomato (*Solanum lycopersicum* L.).

## Materials and methods

### 1. Setup of the experiments and plant material

Two experiments were carried out with tomato plants cultivated in a hydroponic system, in two different times and places. The first experiment was conducted in a controlled-environment growth room at the French Associates Institute for Agriculture and Biotechnology of Drylands, Ben-Gurion University of the Negev, Sde Boker, Israel (online resource 1). The environmental conditions were: a photoperiod of 12/12 hours light/dark, respectively, a relative humidity of 50 %, a temperature of 25 °C, and a light intensity of 200 μmol·m^-2^·s^-1^. Two separated assays were performed under these conditions with tomato plants (*S. lycopersicum*) cv. Ikram, from February-March and April-May, 2016, each time for six weeks, with eight plants per treatment and per control at each cycle (sixteen in total). The plants were not allowed to set fruits due to space limitations.

The second experiment was carried out in a 125-m^2^ mesh house at Universidad del Valle in Cali, Valle del Cauca department, Colombia (online resource 1). This experimental site is located at 967 meters above the sea level, in a sub-humid tropical environment according to Holdridge life zones, with an annual average temperature of 23 °C and a relative humidity of 65 %. Photoperiod and light intensity during the time of this experiment were close to 12/12 hours light/dark and 1100 μmol·m^-2^·s^-1^ in clear days, respectively. Two separated assays were carried out using tomato plants (*S. lycopersicum*) cv. Santa Clara, during April-July and October-December 2017, for twelve weeks each time, with ten plants per treatment and per control at each cycle (twenty in total). The plants were allowed to set fruit until the end of the experiment.

Both experiments (both at Sde Boker and at Cali) consisted of two identical, deep-water hydroponic systems with permanent recirculation of water in 10-liter pots. The pots were connected in a loop with 5-mm diameter tubing to a central pot in which an immersed water pump continuously recirculated the water, at a rate of 1.92 L·min^-1^ (Fig. 1). The MTW system was coupled to a Quantum Biotek magnet, while the control system had a non-magnetized PVC joint. The nutritive solution was Hoagland 50 % solution, adjusted to an initial pH of 6.8 and an electrical conductivity (EC) of 1100 μS·cm^-1^. Additionally, an air bubble supply was introduced into each pot in order to ensure aerobic conditions for the roots. Tomato plantlets were carefully selected from a germination tray, and their roots were thoroughly washed in order to remove any soil residue. The plantlets were attached to a hole in the lid of the pots with foam, so that the solution level covered ¾ of the roots. Every four days, the volume of the solution was reestablished according to the losses from evapotranspiration, in order to maintain the same level.

**Fig. 1.**
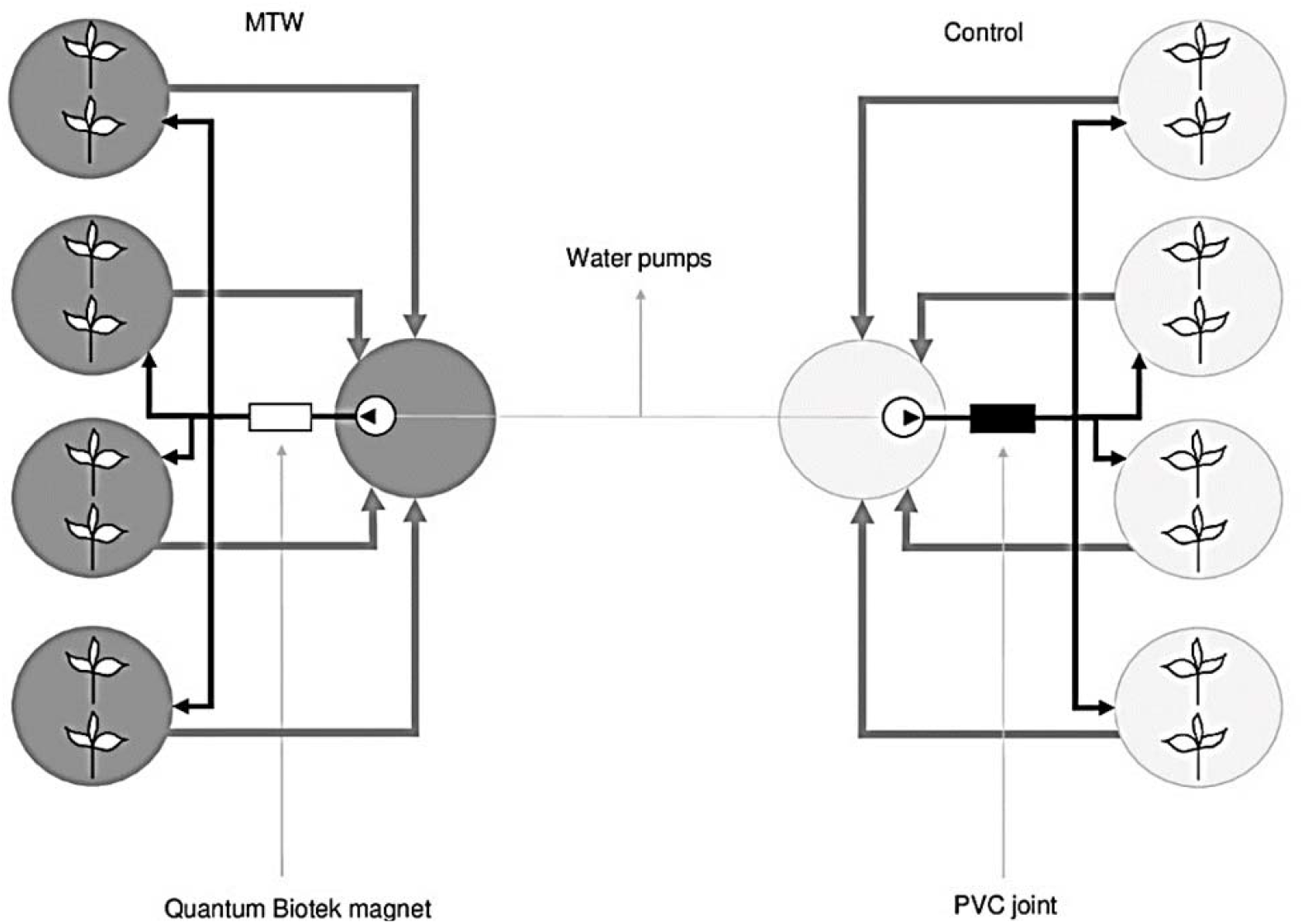
Scheme of the hydroponic system. Arrows show the loop water flow through the magnet (MTW) and the through the non-magnetized PVC joint (control)

### 2. Magnetic treatment device

The Quantum Biotek (Omni Environmental Group, Australia) magnetic device for water treatment comprises a polycarbonate cylinder of 12 cm in length and 2.54 cm in internal diameter. The permanent magnetic field that it generates varies from 0 to 156 mT (millitesla), both in the longitudinal and in the cylindrical direction. The N-S polarity of the magnetic field also varies in both directions. Hence, the water is passed through a non-uniform magnetic field all along the device. For further details of this appliance, see Grewal and Maheshwari (2011).

### 3. Parameters tested in experiment 1 at Sde Boker

#### 3.1. Chlorophyll and gas exchange measurements

Chlorophyll content was estimated with a CCM-200 plus Chlorophyll Content Meter (Opti-Sciences, USA), in relative SPAD units. Gas exchange measurements, including CO_2_ net assimilation rate (*A*), transpiration (*E*) and stomatal conductance to H_2_O (*g_s_*) were estimated at photosynthetic active radiations (PAR) of 850 μmol·m^-2^·s^-1^ and 200 μmol·m^-2^·s^-1^, setting the carbon dioxide concentration at 400 mg·L, the leaf temperature at 25 °C and the relative humidity of the chamber at 38 %, using a LI 6400 Portable Photosynthesis System (LI-COR, USA). The water use efficiency (WUE) was calculated as the ratio between *A* and *E*. Measurements were taken between 10:00 and 15:00. All the latter measurements were done once a week for five weeks for each assay, starting eight days from the setting of the plants in the hydroponic system, choosing the first fully expanded young leaves.

#### 3.2. Water status measurements

For leaf water potential (Ψ_leaf_), the third fully expanded leaves were covered with a moistened plastic bag and carefully detached from the base of the petiole, then rapidly set in an Arimad-3000 Scholander-type pressure chamber (MRCLAB, Israel). This measurement was done three times in each assay, starting two weeks from the setting of the plants in the hydroponic system. The root hydraulic conductance (K_r_) was measured at the end of the assays, in the six-week-old plants, cut at the base of the stem, using a High Pressure Flow Meter (HPFM) (Dynamax, USA) previously calibrated for flow and pressure according to the diameter of the stem.

#### 3.3. Stomatal density, final biomass and C/N composition

Leaf imprints of mature leaves, of both the adaxial and abaxial sides, were taken by covering the surface with silicone using a suitable dispenser. After drying, the peeled imprints were overlaid with nail polish. The films obtained were examined with a microscope at 40X to count the stomata in two sections of the same area (20 μm x 30 μm). At the end of the assays (six weeks), plants were separated and weighed for root, stem and leaf fresh weight. After drying in an oven at 65 °C for 72 hours, dry weight was recorded. For carbon and nitrogen analysis, dry leaves and root samples were ground to powder, then processed in a Flash 2000 Elemental Analyzer (Thermo Fisher Scientific, Netherlands), using 7 samples for treatment and 7 for control.

#### 3.4. Physicochemical measurements of MTW

Two parameters of water were measured (pH and electric conductivity). For pH and electric conductivity, a potentiometer Mettler Toledo and a conductimeter Thermo Scientific device were used, respectively, sampling the water once a week in both control an MTW hydroponic systems, during the time of each assay (six weeks).

### 4. Parameters tested in experiment 2 at Cali

#### 4.1. Fruit harvest and final biomass

Fruits were collected from both treated and untreated plants as ripened fruits stop growing and become partially red. At each assay, five harvests were done until the wilting of the plants; the fruits were counted and weighed in fresh and then oven-dried for 72 hours. Once the plants start to wilt around twelve weeks from the onset of the assays, they were harvested and separated in roots, stems and leaves. After drying in an oven at 65 °C for 72 hours, dry weight was recorded.

#### 4.2. Surface tension of water

The surface tension was assessed by the method of the ring. The ring was hanged to a dynamometer, connected to an interface (LD Didactics, Germany), which displays the readings in a computer graph (Fig. 2). The water samples were first collected from both MTW and control hydroponic systems before starting the recirculation, in order to set a base line of surface tension of the water. Later, water samples were collected at the fifth week from the start of the recirculation, controlling the temperature at 25°C. Each water sample was placed in a beaker on an adjustable base touching the ring, in order to immerse it and obtain the maximum tension readings several times, in mN·m (millinewton per meter). Non recirculated water was used as negative control.

**Fig. 2.**
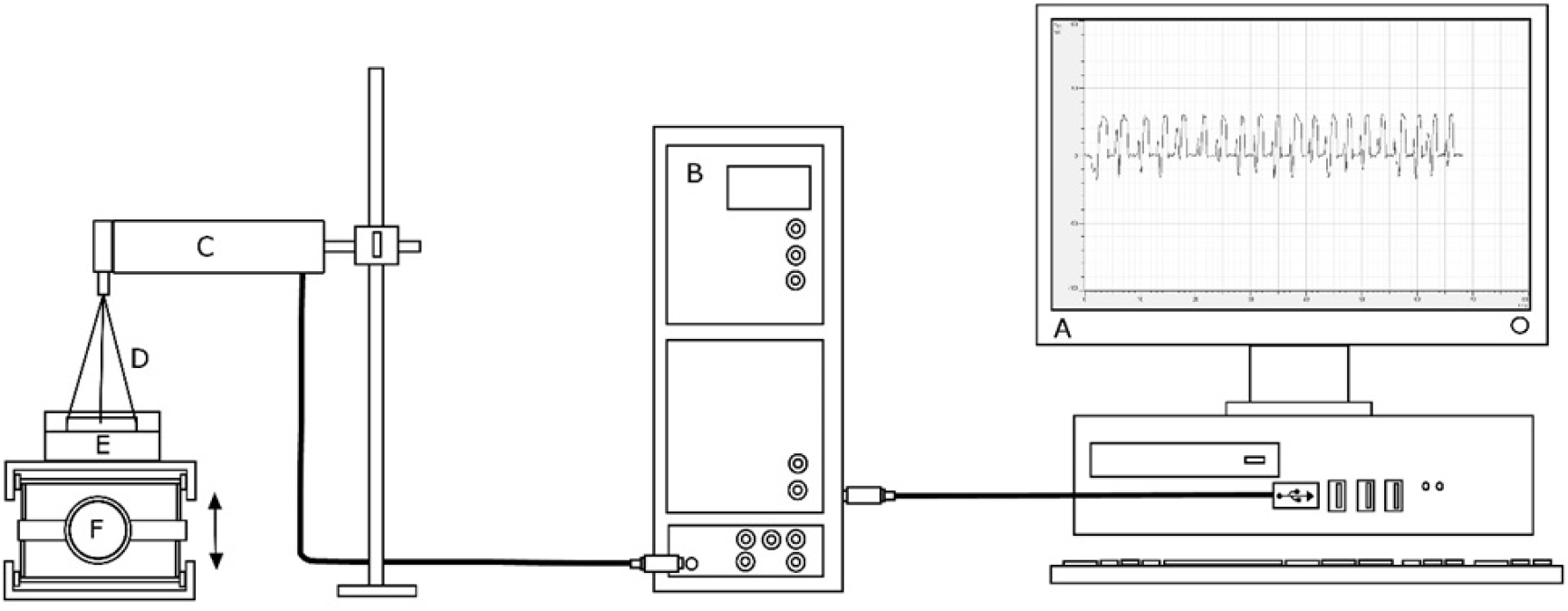
Arrangement of the surface tension measurement. A: computer. B: interface. C: dynamometer. D: pending ring. E: water sample. F: adjustable base

### 5. Statistical analysis

The experiments consisted in a completely randomized design. In the treatment, the Hoagland solution passed through the magnetic device (MTW), whereas in the control, the solution passed through a non-magnetized PVC joint. The analyses were performed using 16 repetitions (eight from the first assay plus eight from the second assay) per treatment and control (an individual plant was considered one repetition), in experiment 1. In experiment 2, there were 20 repetitions (ten from the first assay plus ten from the second assay) per treatment and control. Data for both experiments was processed with IBM SPSS Statistics 22, previously running a test of homogeneity of variances (Levene’s test) and normality (Kolmogorov-Smirnov test). Being that in both tests the null hypothesis of homogeneity of variance and normality in the two groups was not rejected (*P*>0.05), a general linear model procedure for repeated measurements, with a significance of 95 %, was run, testing the significance between and within subjects. For one single measurements (like final biomass) a double-tailed T-test was done.

## Results

### Experiment 1

In relation to the gas exchange parameters, the plants under MTW exhibited higher *A* at the high light intensity evaluated (PAR 850 μmol·m^-2^·s^-1^), with average values of 12.53 μmol CO_2_·m^-2^·s^-1^ for MTW and 11.66 μmol CO_2_·m^-2^·s^-1^ for the control (an increment of 7.46 %), being this effect significant (*P = 0.032*). As for *E*, the response was similar to that of *A*, having an increment of 9.46 % at the higher PAR, being this value also significant (*P = 0.042*). The latter was consistent with a greater *g_s_* of treated plants (10.77 % increment over the control plants), but at a lower level of significance (*P = 0.112*). Despite the increase in transpiration, the WUE of the treated plants was almost the same of that of the control plants. However, when assessing the gas exchange parameters at a low light intensity (PAR 200 μmol·m^-2^·s^-1^), only *E* and *g_s_* kept a higher average in the magnetic treatment, whereas *A* and WUE exhibited lower values than the control plants (although all of the latter values not significant). The chlorophyll content in young leaves was found to be significantly higher (*P = 0.035*) in the plants under MTW than in the control plants (46.96 SPAD units for the treatment vs. 44.34 SPAD units for the control; raise of 5.90 %). Data in Table 1.

**Table 1.** Effect of MTW on the gas exchange parameters and leaf chlorophyll content in hydroponically-grown tomato, according to two light intensities: *A*: photosynthetic rate. *E*: transpiration. *g_s_*: stomatal conductance. WUE: water use efficiency. Chl: chlorophyll content. SE: standard error. Data are means of 16 plants per treatment and control

| Photosynthetically active radiation (PAR) |  |  |  |  |  |  |  |  |
| --- | --- | --- | --- | --- | --- | --- | --- | --- |
| Parameter | 850 $\mu\text{mol}\cdot\text{m}^{-2}\cdot\text{s}^{-1}$ | | | | 200 $\mu\text{mol}\cdot\text{m}^{-2}\cdot\text{s}^{-1}$ | | | |
|  | Treatments | Average | SE | Sig | Treatments | Average | SE | Sig |
| *A* ($\mu\text{mol CO}_2\cdot\text{m}^{-2}\cdot\text{s}^{-1}$) | MTW | 12.53 | 0.29 | 0.032* | MTW | 8.97 | 0.46 | 0.878 |
|  | Control | 11.66 | 0.28 |  | Control | 9.07 | 0.47 |  |
| *E* ($\text{mmol H}_2\text{O}\cdot\text{m}^{-2}\cdot\text{s}^{-1}$) | MTW | 4.05 | 0.14 | 0.042* | MTW | 3.92 | 0.27 | 0.358 |
|  | Control | 3.70 | 0.14 |  | Control | 3.56 | 0.27 |  |
| $g_s$ ($\text{mol H}_2\text{O}\cdot\text{m}^{-2}\cdot\text{s}^{-1}$) | MTW | 0.216 | 0.011 | 0.112 | MTW | 0.25 | 0.02 | 0.396 |
|  | Control | 0.195 | 0.010 |  | Control | 0.22 | 0.02 |  |
| WUE ($\mu\text{mol CO}_2/\text{mmol H}_2\text{O}$) | MTW | 3.32 | 0.09 | 0.681 | MTW | 2.48 | 0.13 | 0.095 |
|  | Control | 3.37 | 0.08 |  | Control | 2.87 | 0.19 |  |
| Chl (SPAD units) | MTW | 46.96 | 0.85 | 0.035* |  |  |  |  |
|  | Control | 44.34 | 0.85 |  |  |  |  |  |

(\*) indicates significant ( $P < 0.05$ ) differences between treatment and control.

Regarding water status parameters, under MTW, tomato plants showed a significantly higher (*P = 0.006*) Ψ_leaf_, reaching average values of −0.33 MPa, which is 13.15 % more than that of the control (−0.38 MPa). Along with this, the K_r_ of the MTW-plants experienced an increase of 74.4 % (1.31 x 10^-7^ m·s^-1^·MPa^-1^; *P = 0.009*) over their control counterparts (7.52 x 10^-8^ m·s^-1^·MPa^-1^) (Fig. 3). The EC in the 50 % Hoagland solution showed a decrease in the system with no magnetic treatment. This means that the solution passing across the magnetic device maintained a higher EC during the evaluation period. In contrast, the average pH was found to be slightly lower in the magnetically-treated Hoagland solution (Figure 3).

**Fig. 3.**
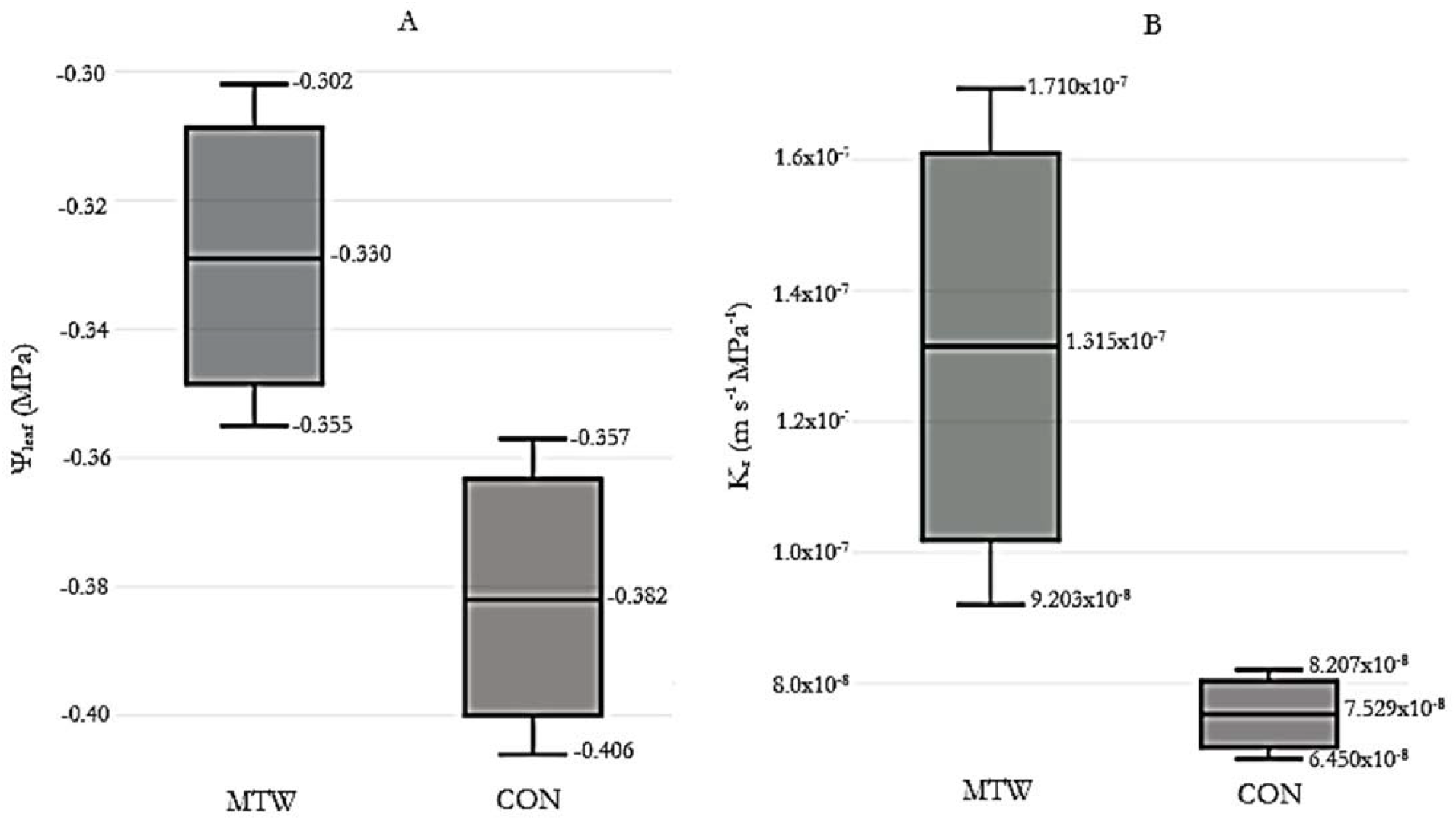
Measurements of A: leaf water potential (*P = 0.006*\*\*) and B: root hydraulic conductance in six-week-old tomato plants (*P = 0.009*\*\*). Middle lines, boxes and whiskers represent the average, 50 % of the data and 95 % confidence interval, respectively, of 16 plants. (**) means significant (*0.001 < P < 0.01*) differences between treatment and control

The difference in stomatal density between the treated and control plants in both the adaxial and abaxial sides of the leaves was found to be not statistically significant, although the number of stomata in the abaxial side of plants under MTW was slightly higher (5.81 %). As for the fresh biomass of the six-week-old plants, the control plants showed less fresh weight in stems and leaves than the treated plants, but regarding dry weight, the values were practically equal (Table 3).

**Table 2.** Stomatal density and biomass accumulation in treated and control plants. Data are means of 16 plants in each treatment

| Stomatal density (stomata/1000 μm<sup>2</sup>) |  |  |  |  |  |  |
| --- | --- | --- | --- | --- | --- | --- |
|  |  | Average | Min | Max | SD | Sig |
| Abaxial | MTW | 27.28 | 15 | 51 | 7.88 | 0.413 |
|  | Control | 25.78 | 14 | 41 | 6.62 |  |
| Adaxial | MTW | 6.81 | 1 | 16 | 3.15 | 0.825 |
|  | Control | 6.97 | 2 | 13 | 2.31 |  |
| Fresh weight (g/plant) |  |  |  |  |  |  |
|  |  | Average | Min | Max | SD | Sig |
| Roots | MTW | 43.45 | 28.72 | 57.21 | 9.52 | 0.824 |
|  | Control | 44.17 | 24.79 | 52.58 | 8.47 |  |
| Stems | MTW | 84.73 | 42.61 | 139.75 | 29.12 | 0.746 |
|  | Control | 81.71 | 39.50 | 126.72 | 22.02 |  |
| Leaves | MTW | 175.26 | 85.70 | 349.99 | 65.72 | 0.684 |
|  | Control | 166.96 | 95.51 | 248.38 | 45.50 |  |
| Dry weight (g/plant) |  |  |  |  |  |  |
|  |  | Average | Min | Max | SD | Sig |
| Roots | MTW | 2.77 | 1.90 | 3.54 | 0.544 | 0.706 |
|  | Control | 2.84 | 1.76 | 3.40 | 0.517 |  |
| Stems | MTW | 6.31 | 2.54 | 11.37 | 2.70 | 0.929 |
|  | Control | 6.38 | 3.03 | 9.57 | 1.87 |  |
| Leaves | MTW | 16.65 | 7.04 | 31.78 | 7.00 | 0.810 |
|  | Control | 16.15 | 8.40 | 23.91 | 4.32 |  |

The C and N composition in the dry tissues varied in relation to the C/N ratio, with the plants under MTW having a higher ratio (*P = 0.045*) in the roots (11.16) than the control plants (10.06). This was derived from the lower N content in the roots of the treated plants, because the C content was statistically equal in both root and leaves. Nitrogen content in leaves was slightly lower in MTW-plants but not at a significant level (Table 3).

**Table 3.** Carbon and nitrogen composition of roots and leaves of tomato plants under MTW. Data are means of 16 plants. (*) indicates significant (*P < 0.05*) differences between treatment and control according to t-test.

| Elemental composition of roots and leaves (% dry matter) |  |  |  |  |
| --- | --- | --- | --- | --- |
|  | Treatments | Average | SE | Sig |
| C content roots | MTW | 39.44 | 0.15 | 0.390 |
|  | Control | 39.77 | 0.34 |  |
| C content leaves | MTW | 38.38 | 0.34 | 0.087 |
|  | Control | 39.22 | 0.27 |  |
| N content roots | MTW | 3.54 | 0.07 | 0.017* |
|  | Control | 3.98 | 0.14 |  |
| N content leaves | MTW | 4.69 | 0.21 | 0.689 |
|  | Control | 4.84 | 0.29 |  |
| C/N ratio roots | MTW | 11.17 | 0.21 | 0.045* |
|  | Control | 10.07 | 0.43 |  |
| C/N ratio leaves | MTW | 8.27 | 0.39 | 0.980 |
|  | Control | 8.26 | 0.49 |  |

### Experiment 2

The two assays carried out in the mesh house showed a consistent pattern of tomato plants growing in the hydroponic-MTW system having higher biomass accumulation in all organs (roots, stems, leaves and fruits) in comparison with control plants. The latter effect was significant (*P < 0.05*) in fresh weight of fruits per plant and number of fruits per plant (Figures 3 and 4), although not for dry weight of fruits. The relative increment in the previous parameters were of 61.7 % for fresh weight of fruits, 39.9 % for dry weight and 85.3 % in number of fruits in MTW-grown plants over their control counterparts. From this outcomes, it can be inferred that, despite the large increase in fruit number per plant in MTW, the average dry biomass of each fruit was lower than in control plants, which explains the lower increment in this parameter and its lower level of significance. Relative to the biomass of roots, stems and leaves, the increment of dry weight was of 30.3 % for roots, 13.6 % for stems, and 10.1 % for leaves, yielding a total increase in dry biomass of 12.6 % (not considering the dry biomass of fruits). Notwithstanding, it was only found a significant difference in root dry weight.

**Fig. 4.**
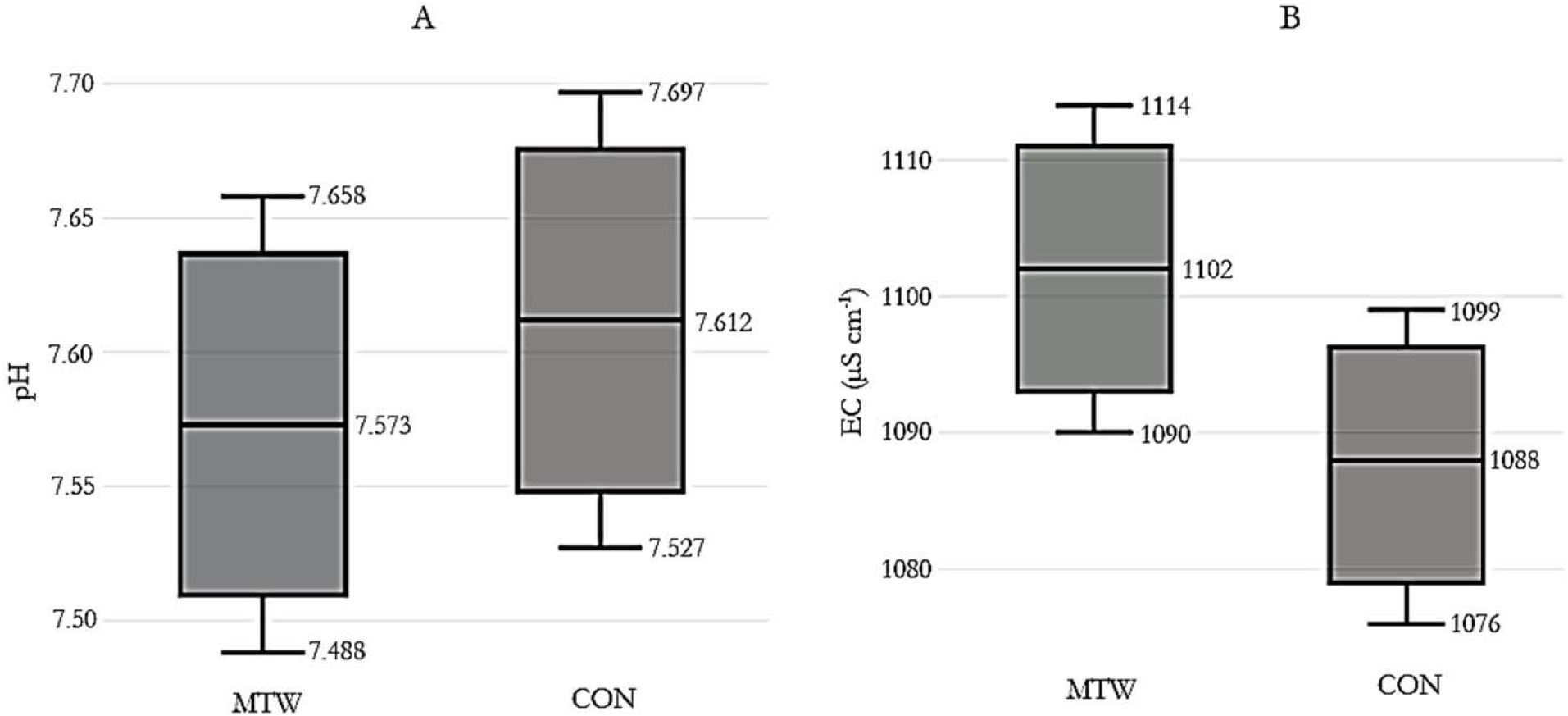
Measurements of A: pH (*P = 0.519*) and B: electric conductivity (EC) (*P = 0.095*). Middle lines, boxes and whiskers represent the average, 50 % of the data and 95 % confidence interval, respectively. Data are means of 30 repeated measurements in the solution

**Fig. 5.**
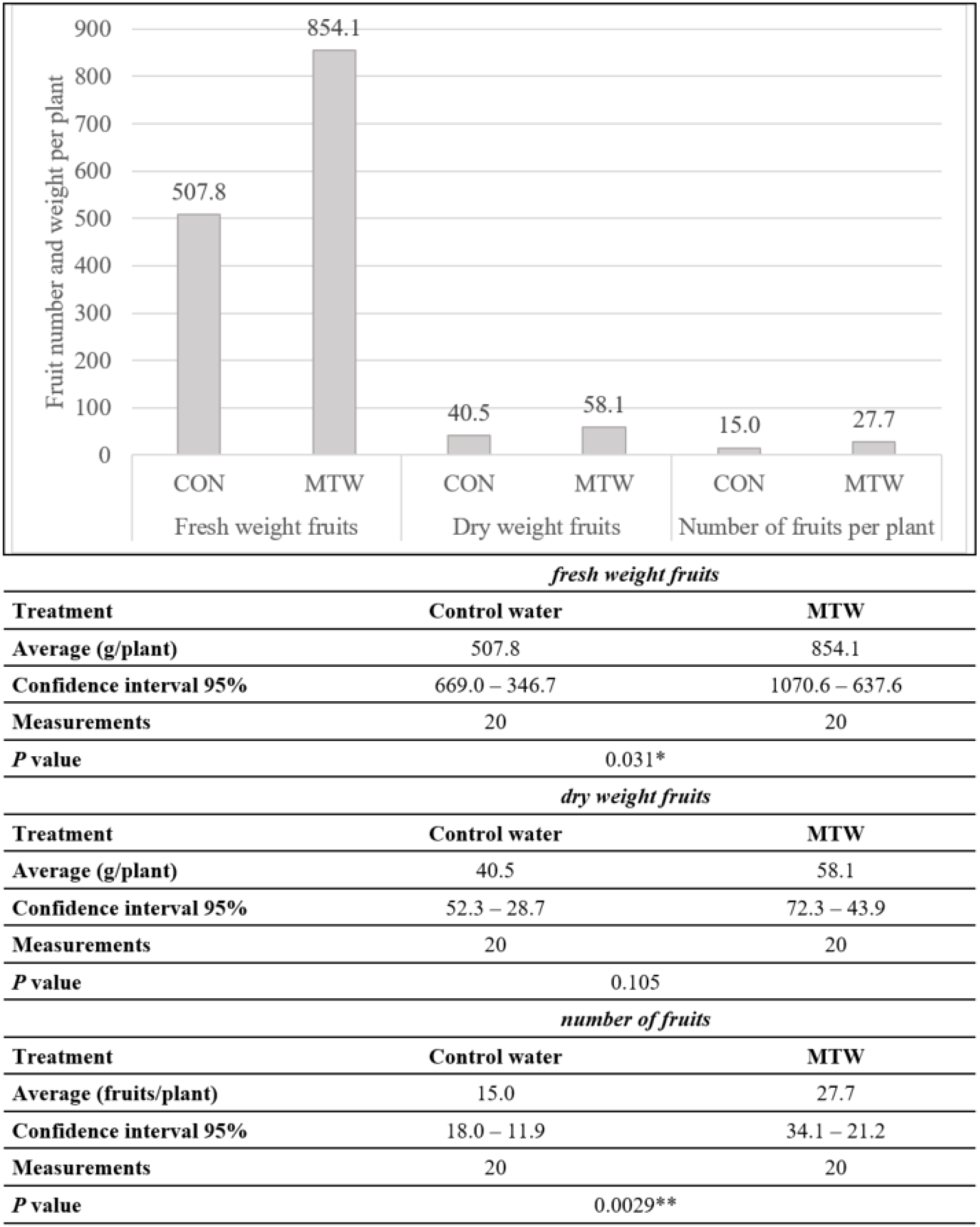
Measurements of fresh weight of fruits, dry weight of fruits and number of fruits per plant. Data are means of 20 plants per treatment and control. (**) means significant (*0.001 < P < 0.01*) differences between treatment and control. (*) means significant (*0.01 < P < 0.05*) differences between treatment and control

**Fig. 6.**
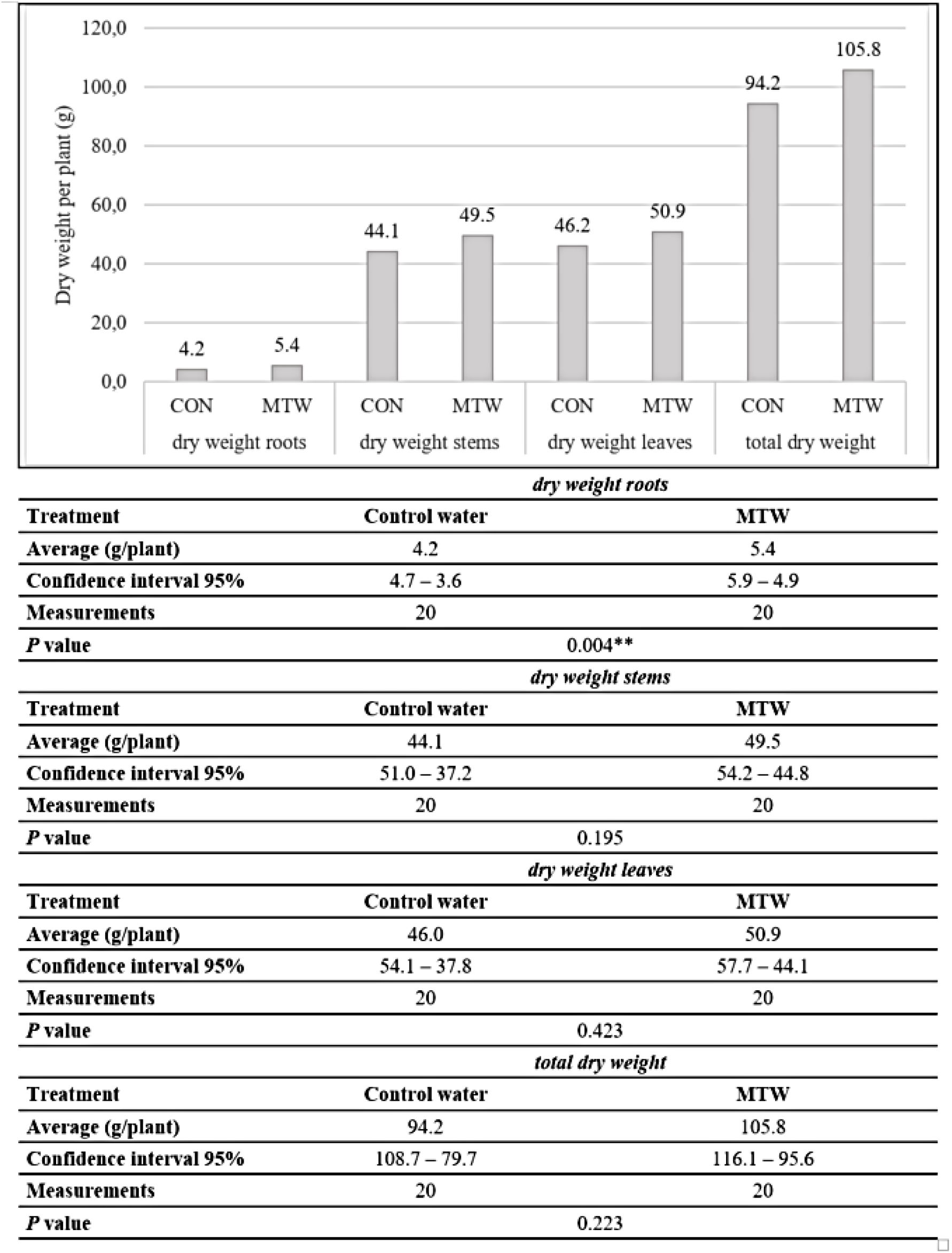
Dry weight of roots, dry weight of stems, dry weight of leaves and total dry weight (not considering fruit weight). Data are means of 20 plants per treatment and control. (**) means significant (*0.001 < P < 0.01*) differences between treatment and control. (*) means significant (*0.01 < P < 0.05*) differences between treatment and control

The analysis of the data of this measurement revealed that the initial surface tension of water with Hoagland solution in both systems was practically the same, because there was not still any magnetic effect; as expected, this water at rest had a higher surface tension than the recirculated water, regardless of the treatment. However, when the recirculation begun and the water passed through the magnetic device, there was a decrease in surface tension of 1.1 % in relation to control water, having a non-significant P-value (*P* = 0.223).

**Fig. 7.**
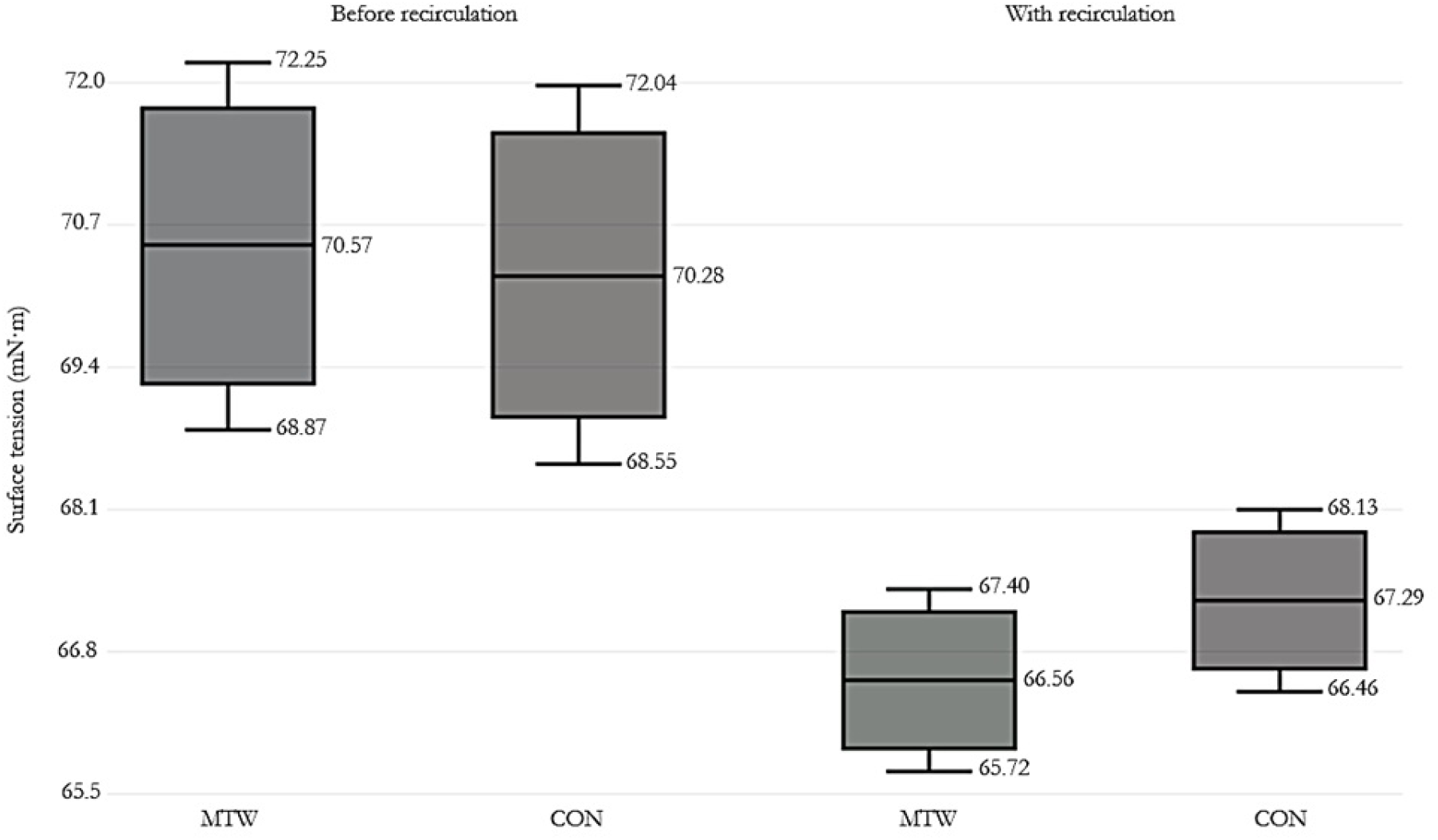
Measurements of surface tension of water with 50 % Hoagland solution. Base line of surface tension of water at rest is at left (*P = 0.824*), and surface tension of running MTW and control water is at right (*P = 0.223*). Middle lines, boxes and whiskers represent the average, 50 % of the data and 95 % confidence interval, respectively. Data are means of 229 and 232 measurements for MTW and control, respectively, before recirculation, and 1032 and 1020 measurements for MTW and control, respectively, with recirculation

**Table 4.**
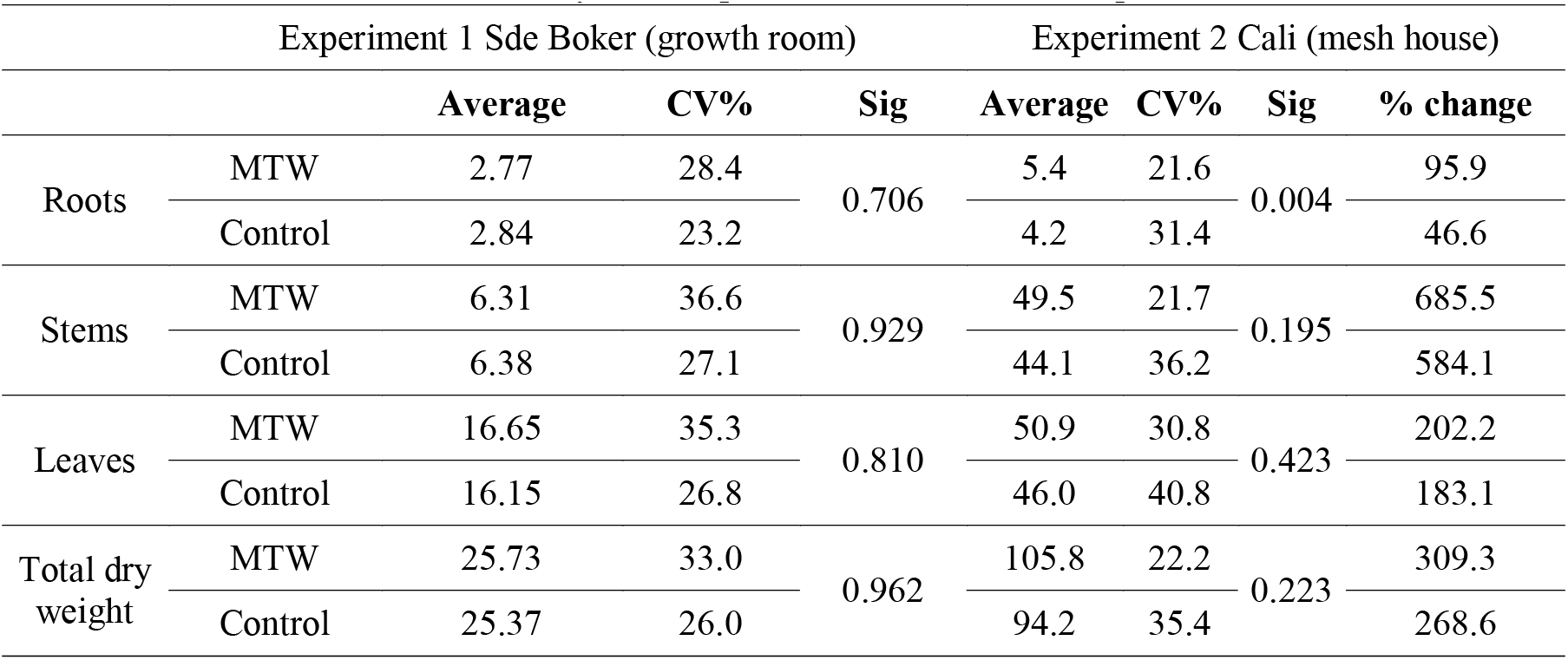
Biomass yield comparison between the two experiments

## Discussion

Several physiological effects in tomato plants were recorded in this work, attributed to the magnetically treated-water, although the specific changes in the physicochemical properties of the water (pH, EC and surface tension) were of arguable significance. Among the physiological effects, there was a significant enhancement in the CO_2_ net assimilation and transpiration rates, chlorophyll content, root hydraulic conductance and leaf water potential. However, the latter effects did not lead to a higher biomass accumulation in the MTW-grown plants in the growth room experiment. This is ascribed the low light conditions of the growth room (200 μmol·m^-2^·s^-1^), since the best performance of photosynthesis was achieved around 850 μmol·m^-2^·s^-1^. Hence, it can be inferred that the significant increase in biomass yield in the mesh house experiment were related to the natural light conditions of the site, which enabled the MTW-plants to perform at greater photosynthetic rates, which lead to higher CO_2_ assimilation.

The latter is consistent with previous reports about tomato yield in hydroponic systems being higher when using a light intensity similar to that of spring than with that similar to winter light conditions (Logendra et al., 2001). Other authors report that, when growing hydroponic tomato under greenhouse production, the application of supplemental light on the canopy of the plants increases biomass accumulation, when light insufficiency is a limiting factor (Jiang et al., 2017). In addition, the enhancing effect of MTW on biomass yield (at experiment 2) was also consistent with other reports where this treatment elicited a significant increase in fruit yield, even at limited water or nutrient supply but in presence of full sunlight (Yusuf et al., 2017a; Ogunlela and Yusuf, 2016; El-Yazied et al., 2012). The previous trend indicates that, in adequate light conditions for tomato plant development, the application of MTW results in a positive yield effect.

Our results regarding the photosynthetic response to MTW at high light intensities are consistent with those reported by Moussa (2011). This author also found a significant increase in the photosynthetic activity and total chlorophyll content in beans planted in a growth room on a sandy-clayey substrate, with similar light intensity, temperature and humidity conditions to this experiment. Likewise, Marei et al. (2014) observed an increase in the chlorophyll content in bell pepper irrigated with magnetically-treated brackish water, in an open-field experiment. They attributed this effect to the high nutrient absorption from the treated water through the roots, compared to the untreated water. About leaf chlorophyll content, this is a good predictor of productivity, because it is strongly correlated with the photosynthetic processes (Ciganda et al., 2009). However, in this work, we do not necessarily ascribe the higher CO_2_ assimilation to the greater chlorophyll content observed in the treated plants, since the stomatal effects on photosynthesis in tomato can be stronger than the chlorophyll effects (Romero-Aranda et al., 2001).This might indicate that the metabolic water that has passed through the magnetic field is exerting a stomatal effect, a process associated with the transpiration rate, the root hydraulic conductance and the leaf water potential (Garcia et al., 2007).

Modification of *g_s_* within a species through changes in stomatal density is a process that depends on the ecophysiological conditions where the plant is settled; for example, in tomato plants growing in a saline substrate or under a water stress regime, the stomatal density was strongly reduced (Romero-Aranda et al., 2001; Sam et al., 2000). Stomatal density is positively correlated with *g_s_*, *A* and WUE (Xu and Zhou, 2008). From our results, it can be inferred that although MTW produced only a slight effect on stomatal density (in the abaxial side of the leaves), this might have contributed to the increased *g_s_* observed in the treated plants. Here, we also observed an augmentation in stomatal conductance in the MTW-grown plants (although the *P* value was only below 11.2 %); this phenomenon was also observed by other authors in cowpea grown in a soil substrate and subjected to MTW (Sadeghipour and Aghaei, 2013). Stomatal conductance/resistance is a trait highly sensitive to soil and atmospheric water deficit, and it can severely restrict the CO_2_ uptake and the release of H_2_O in the process. From the latter arguments, it is likely that a greater *g_s_* in the MTW-grown plants in the mesh house experiment was the cause of the significant increase in the number of fruits and in the dry biomass. On the other hand, since the plants in the growth room experiment were grown in full moisture, under non-stressing conditions, the only factor that could have affected *g_s_* was the magnetically-treated water. The augmentation of both CO_2_ and H_2_O exchange explains why WUE was not altered in the plants subjected to MTW.

The above mentioned is consistent with the fact that an increase in Ψ_leaf_ leads to higher *g_s_* and *E*, both phenomena observed in tomato under MTW in this experiment. It has been comprehensively reported that Ψ_leaf_ is directly related to *g_s_* and *E* in tomato and other species (Assis Gomes et al., 2004; Else et al., 2001). Hence, it is proposed that the mechanism of MTW is more a type of water transport, since the incremental trend of K_r_ and *E* shows that the water transport along the plant to the atmosphere is enhanced. The relationship between *E*, Ψ_leaf_ and K_r_ has been found to be almost directly linear in several crops under flooding, drought and salinity stress conditions (Navarro et al., 2007; Trifilo et al., 2003; Else et al., 2001), so that *E* and Ψ_leaf_ increase or decrease when K_r_ does as well, as a result of the strengthening or weakening of such stressors. One should not underestimate the hydraulic signaling that controls *g_s_*; although abscisic acid (ABA) has been described as the essential elicitor of stomatal closure, transpiration rate and hydraulic conductance are also strongly correlated to this response, being that the water flux must be equal to both the losses through the stomata and the liquid phase transport, and given that several experiments have shown a stomatal closure independent from Ψ_leaf_ but very consistent with a hydraulic signaling via reduced hydraulic conductance (Comstock, 2002).

A high and significant root hydraulic conductance was observed in MTW-grown plants in the growth chamber experiment, as well as higher transpiration in leaves. Here, it is important to highlight that other authors have also found an increase in evapotranspiration in tomato plants when using MTW for irrigation, in addition with a reduction in the viscosity of the treated water (Yusuf and Ogunlela, 2017b). These response is ascribed to a higher water absorption from the plants, which eventually increased the rate of vegetative growth of the tomato plants (Yusuf and Ogunlela, 2017b). Water evaporation from the boundary layer also works as a motion force that pulls the water column, thanks to the strong hydrogen bonds between water molecules; the latter is linked to the root hydraulic conductance. Several studies show that water exposure to magnetic fields enhances the evaporation rate of water. Results from Guo et al. (2012) showed that under a magnetic field of at least 8 tesla, this parameter increases significantly, a trend that was also reported by Holysz et al. (2007). Other authors had indicated that under the influence of a magnetic field, the structure of the liquid water changes and more water molecules are forced between the water shells. These molecules connect the shells and hence create a more stable water-water network (Chang and Weng, 2006). Therefore, stronger hydrogen bonds and weaker stress interactions in the water column along the xylem vessels, is consistent with a looser water transport.

Physical properties of water are involved in xylem water transport; this is a process in which surface tension plays a critical role, helping to create a negative pressure in the water-air interface at the intercellular spaces beneath the substomatal chamber (Domec, 2011). Ultimately, the surface tension generated at the air/water interfaces of cell walls is assumed to be transmitted through a continuous water column to the roots (Kim et al., 2014). Surface tension in the xylem’s sap has such a role, that there is evidence of surfactant substances in the xylem of several angiosperm species, whose function are thought to be related with supporting water transport under negative pressure (Schenk et al., 2017), although it seems contradictory that these surfactant substances may facilitate the formation of bubbles in the xylem, which would actually impair water transport. Nevertheless, the evidence show that xylem surfactants support water transport under negative pressure as explained by the cohesion-tension theory by coating hydrophobic surfaces and nanobubbles, thereby keeping the latter below the critical size at which bubbles would expand to form embolisms (Schenk et al., 2017). Magnetically treated water is also known to exhibit lower surface tension, which altogether with evaporation rate, can generate a greater motion force enhancing water transport through the soil-plant-atmosphere continuum. It is important to take into account that, in this experiment, it was observed a decrease of 1.1 % in the surface tension of MTW. Authors as Cho & Lee (2005) observed a reduction of 8 % in this parameter after passing the water several times through a magnet of 0.16 tesla, an effect that is in accordance to several later reports where magnetically-treated water manifested a reduction in this parameter (Cai et al., 2009; Pang & Deng, 2008). However, it is likely that the method used in this work to measure this property was not accurate enough to reduce the variance and to generate a significant difference between MTW and control.

The reductive trend found in the pH of the Hoagland solution, along with the increasing EC, is consistent with results reported by Grewal and Maheshwari (2011) but opposite to the results of Yusuf and Ogunlela, (2017b). However, while the former authors used the same magnetic device of this experiment, finding a significant decrease in pH and a slight increase in EC, the latter authors used a different device with lower magnetic induction. In this study, the reduction in pH and the increase in EC can be attributed not only to the effect of the magnet, but also to the higher water uptake of the plants, which led to a steady concentration of solutes in the solution.

One feasible reason for the apparent lack of relation between nitrogen tissue content and chlorophyll in the leaves of plants under MTW is that there is not always a clear relationship between these two traits. For example, in winter wheat grown under equal initial N supplements but in contrasting environmental conditions, the correlation between the total N concentration and chlorophyll in the leaves was absent (Shadchina and Dmitrieva, 1995). However, the lower N content in the roots helped to boost the C/N ratio, which indicates a more efficient carbon assimilation per unit of nitrogen. In relation to this, other researchers have found that the content of ash, phosphorus and crude protein in tomato fruits increased when MTW was applied (Han et al., 2004). This may indicate that mineral translocation in the plant and incorporation of major nutrients (as phosphorus and nitrogen) in the tissues can be positively affected by MTW. The findings of El-Yazied et al. (2012) support this theory, in view that they also reported an increase in phosphorus in the leaves.

The magnetic treatment of water for agriculture, although have been documented since the last century (Noran et al., 1996; Lin and Yotvat, 1990), is an issue that still causes some distrusts. This is due to the lack of standardization of the different magnetic treatments, the variability among plant species and the possible interference of the soil or substrate on the magnetically treated-water. Here, these sources of variability are overcoming by using a standard magnetic device in a closed-loop hydroponic system, which assures no loss of magnetic priming over time and distance. This is a fundamental aspect to be considered since several authors mention that water can lose its “magnetic memory” (Teixeira da Siva and Dobranszki, 2014; Otsuka and Ozeki, 2006).

## Conclusions

- MTW exerts an incremental effect on CO_2_ assimilation and H_2_O outflow in tomato plants grown in hydroponic conditions. This was consistent with a stomatal mechanism, either by a wider opening of the guard cells or by an increased stomatal density, although the uncertainty related to the latter parameters was arguable.
- Water transport through the plant was enhanced as a consequence of MTW, due to a coherent relation between greater root hydraulic conductance, less negative leaf water potential and a higher leaf transpiration rate. We propose a surface tension and evaporative mechanism for this effect, since magnetically-treated water is known to exhibit modifications in these properties.
- The reduction in surface tension of MTW observed in this work, although did not reach high levels of significance, is consistent with several previous reports of other authors mentioning this same effect.
- The significant gain in number and fresh biomass of fruits, as well as dry biomass of roots, is likely associated to the above mentioned stomatal and transpiration mechanism, which allowed a higher CO_2_ assimilation. However, this effect was only visible under full sunlight, not under artificial light.
- Although more research is needed in hydroponic conditions, the technology of MTW could be an effective method to improve tomato yield under this production system.

## Supporting information

Experimental setup

Graphical abstract

## Declarations

### Funding

This research did not receive any specific grant from funding agencies in the public, commercial, or not-for-profit sectors.

### Conflicts of interest/Competing interests

The authors declare that there are no conflicts of interest.

### Authors’ contributions

Daniel I. Ospina-Salazar: setup of the experiments, data collection, data analysis, writing

Shimon Rachmilevitch: revision and data analysis

Santiago Cuervo-Jurado: setup of the experiments, data collection Orlando Zúñiga-Escobar: revision and writing

## Acknowledgements

The authors offer thanks to the Administrative Department of Science, Technology and Innovation of Colombia (COLCIENCIAS) and the Jacob Blaustein Institutes for Desert Research for the economic support and the collaboration between the Valle University at Cali-Colombia and the Ben-Gurion University of the Negev at Sde Boker, Israel.

## Notes

### Competing Interest Statement

The authors have declared no competing interest.

