## Supplementary material for "Biomass accumulation and physiological responses of tomato plants to magnetically–treated water in hydroponic conditions": Experimental setup

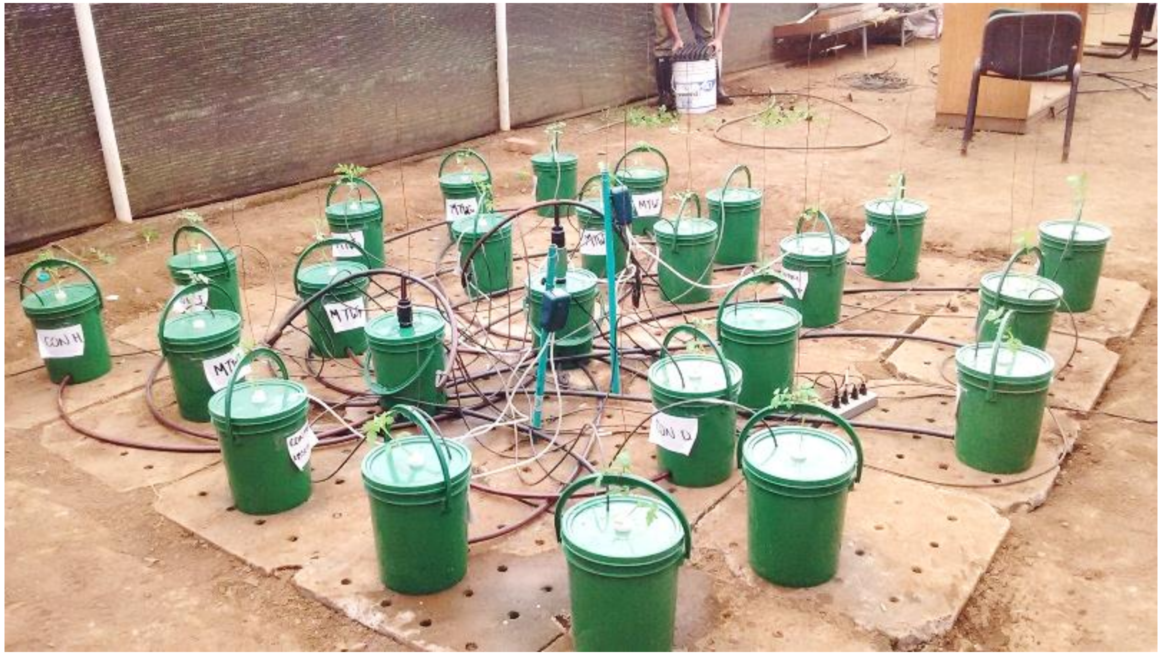

View of the experiment at Cali

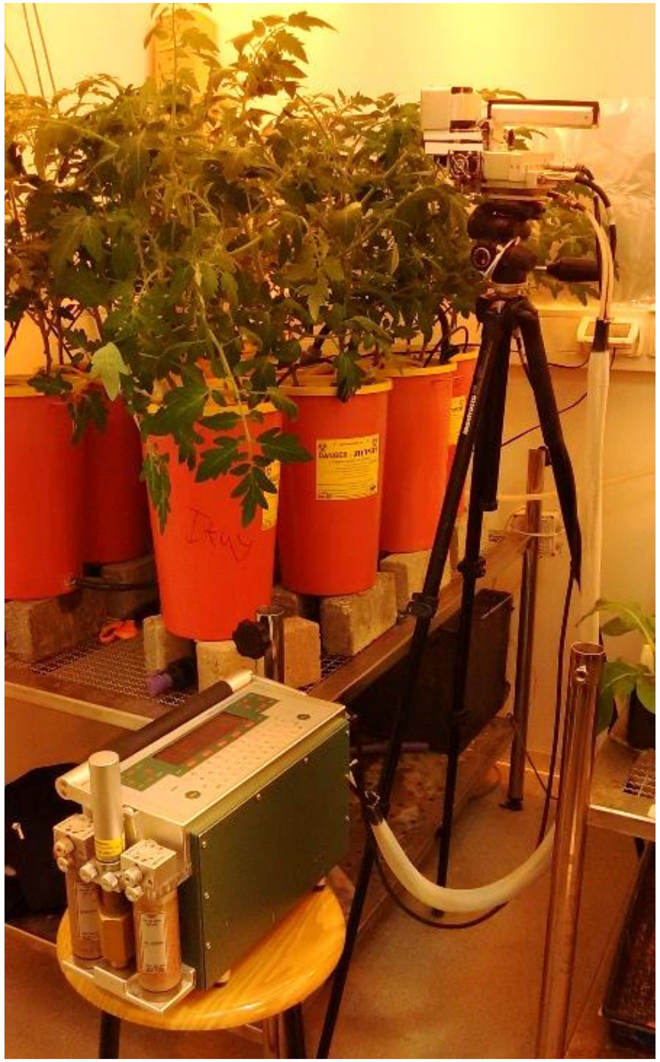

View of the experiment at Sde Boker, measuring photosynthesis with LICOR-6400
