## Supplementary material for "Biomass accumulation and physiological responses of tomato plants to magnetically–treated water in hydroponic conditions": Graphical abstract

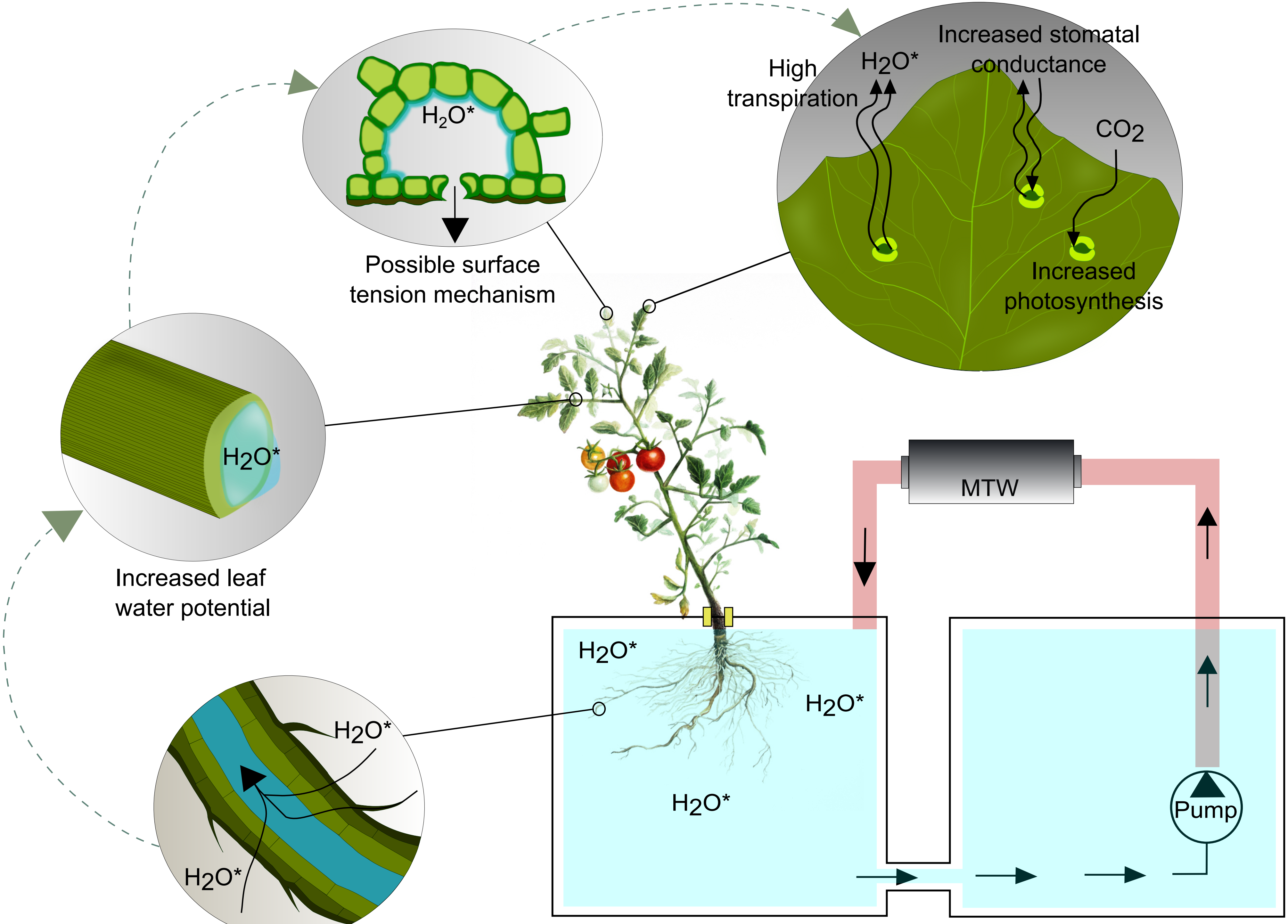


$H_2O^*$: Magnetically-treated water molecule
MTW: Magnetic treatment of water

Copyright: Ospina-Salazar & Benavides, 2016
